# Morphological comparison of astrocytes in the lamina cribrosa and glial lamina

**DOI:** 10.1101/2024.09.07.610493

**Authors:** Susannah Waxman, Hannah Schilpp, Ashley Linton, Tatjana C. Jakobs, Ian A. Sigal

## Abstract

**Purpose:** Although the mechanisms underlying glaucomatous neurodegeneration are not yet well understood, cellular and small animal models suggest that LC astrocytes undergo early morphologic and functional changes, indicating their role as early responders to glaucomatous stress. These models, however, lack the LC found in larger animals and humans, leaving the *in situ* morphology of LC astrocytes and their role in glaucoma initiation underexplored. In this work, we aimed to characterize the morphology of LC astrocytes *in situ* and determine differences and similarities with astrocytes in the mouse glial lamina (GL), the analogous structure in a prominent glaucoma model.

**Methods:** Astrocytes in the LCs of twenty-two eyes from goats, sheep, and pigs were stochastically labeled via Multicolor DiOlistics and imaged *in situ* using confocal microscopy. 3D models of DiOlistically-labeled LC astrocytes and hGFAPpr-GFP mouse GL astrocytes were constructed to quantify morphological features related to astrocyte functions. LC and GL astrocyte cross-pore contacts, branching complexity, branch tortuosity, and cell and branch span were compared.

**Results:** LC astrocytes displayed distinct spatial relationships with collagen, greater branching complexity, and higher branch tortuosity compared to GL astrocytes. Despite substantial differences in their anatomical environments, LC and GL astrocytes had similar cell and branch spans.

**Conclusions:** Astrocyte morphology in the LC was characterized through Multicolor DiOlistic labeling. LC and GL astrocytes have both distinct and shared morphological features. Further research is needed to understand the potentially unique roles of LC astrocytes in glaucoma initiation and progression.

## Introduction

The lamina cribrosa (LC) is an initial site of injury to the retinal ganglion cell axons in glaucoma.^1,2^ This injury is known to result in the progressive and irreversible optic neuropathy characteristic of the disease. The underlying mechanisms causing glaucomatous neurodegeneration are poorly understood. Intraocular pressure (IOP) is often elevated in glaucoma. IOP is currently the only clinically modifiable risk factor in treatment of the disease. However, even with pharmacotherapies and surgeries aimed to control pressure, many patients continue to suffer from progressive optic neuropathy and as a result, progressive vision loss. This points to a clear need to better understand the basic *structure and function* of the LC, and particularly its components influencing disease initiation and progression.

In health, astrocytes are known to play critical roles in supporting neurons through processes including signaling across networks,^3–5^ neurovascular coupling,^6,7^ and responding to local mechanical strain.^8–10^ In the LC, astrocytes are the most abundant glial cell type, accounting for over 90% of cell nuclei in human lamina.^11^ In cellular and animal models of glaucoma, LC astrocytes undergo morphologic and functional changes early in the process of pathogenesis,^12–17^ pointing to astrocytes as early responders to glaucomatous stressors in the LC. Importantly, IOP-induced astrocyte morphologic changes have been detectable before any observable neurodegeneration,^13,18,19^ There has been long-standing interest in better understanding the morphology of these astrocytes and its relation to their roles in health and glaucoma. Previous work has relied largely on in vitro models, mouse models, and large animal or human donor samples. The unique benefits and important limitations of these previous works are discussed below.

Isolation and culture of LC astrocytes *in vitro*^20,21^ has allowed excellent access and experimental flexibility for investigation of these cells^22–24^. A limitation of these in vitro models is that before evaluation of astrocytes, they are removed from their native tissue environment. This impacts the inputs they normally receive as part of 3D tissue under complex and variable mechanical strain. Culture of cells on flat and/or unphysiologically stiff materials such as conventional polystyrene plates is known to alter cell morphology.^25,26^ Additionally, isolation and in vitro culture of astrocytes removes the opportunity for interaction with other cell types. In tissues, astrocytes exist in close association with extracellular matrix and other cell types of the neurovascular unit. As both tissue mechanics^27,28^ and cell-cell communication^4,29–31^ are important aspects of LC physiology in health and disease, models that are able to preserve these features are likely to more closely represent what takes place in the eyes of healthy people and glaucoma patients.

A large portion of what is known about the morphology of these astrocytes *in situ* comes from research conducted in mouse models.^13,15^ This work has been able to reveal the detailed structures of astrocytes in the glial lamina (GL), the mouse analog of the collagenous LC that humans and other large mammals have.^32–34^ Mouse models of glaucoma are powerful tools that have allowed investigation of the earliest changes these astrocytes undergo, *in situ*, in response to elevated IOP. Mouse glaucoma models have shown IOP-induced astrocyte branch retraction and branching simplification^30,35,36^ which may result in impaired contact and communication with blood vessels, RGC axons, and neighboring astrocytes. Astrocyte structural changes are likely to alter vital processes necessary for local support to axons. Notably, the degree of IOP-induced astrocyte structural changes correlated with glaucoma severity.^13^

However, GL astrocytes cannot be assumed to faithfully represent LC astrocytes. The LC, between the retina and the optic nerve, is part of the central nervous system. In other central nervous system tissues, mouse astrocytes differ substantially in structure from those of larger mammals, including non-human primates and humans. For example, human cortical astrocytes are significantly larger and more structurally complex than their rodent counterparts, with 10-fold more GFAP+ primary processes^37^. Structural differences were coincident with functional differences, with human astrocytes propagating calcium signals several-fold faster than their rodent counterparts.^37^ Additionally, the organization and abundance of collagen in the LC and GL is distinct (**Fig. 1**.)^38,39^ Astrocytes in the LC exist in close association with the dense network of collagenous beams. This collagen network plays an important role in the mechanics of the tissue and is substantially different in the GL. Given the limited success of many therapies to translate between mouse studies and clinical trials, investigation of models that represent human anatomy and physiology to the best of our ability are critical.

**Figure 1:**
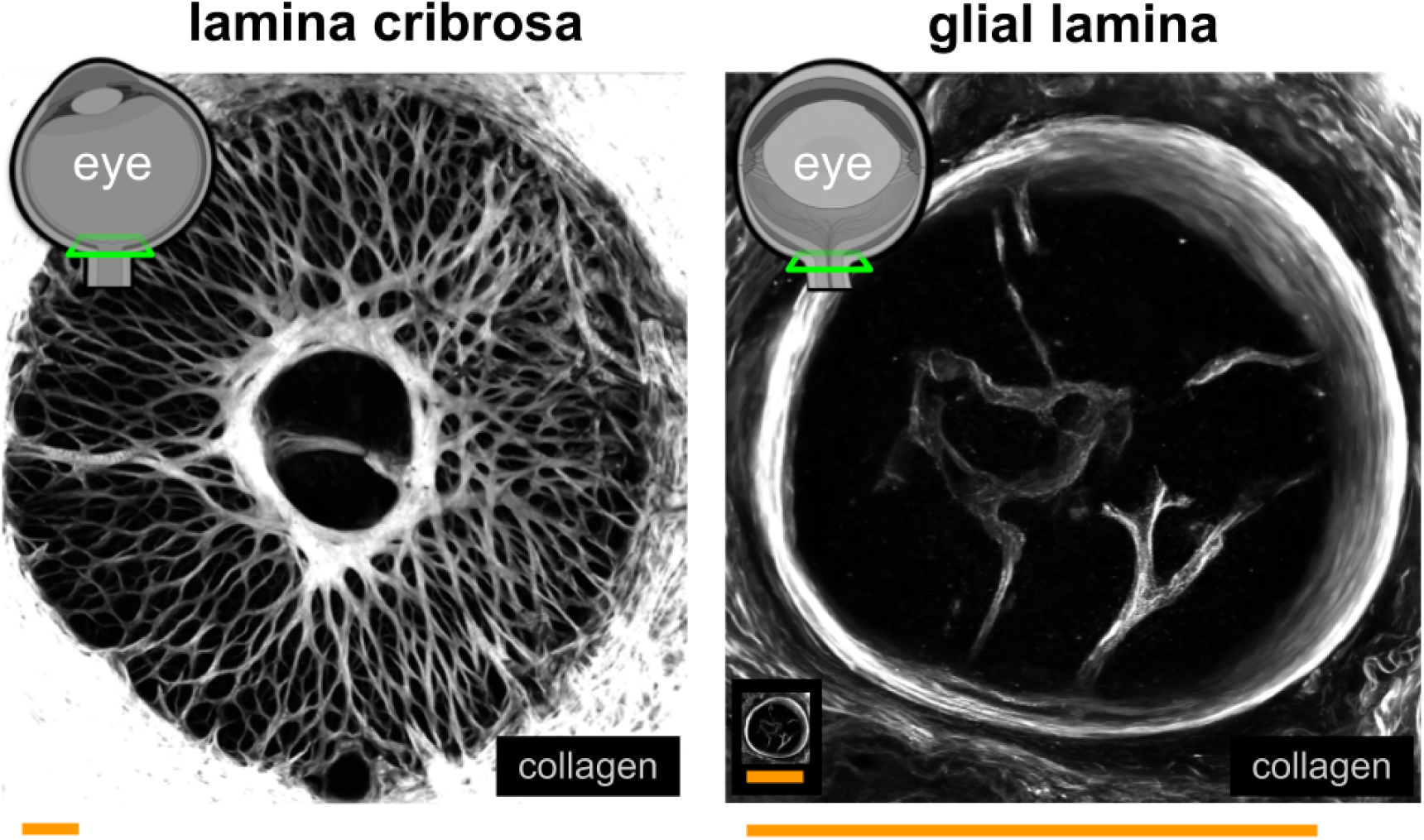
The LC and GL are structurally distinct. Collagen organization and canal size differ substantially among the LC and GL. Example images of the goat LC (left) and mouse GL (right.) Neural tissues (dark regions) in the LC are divided into pores by a network of robust collagenous beams (grayscale.) Neural tissues of the GL are not divided into pores by collagen beams. The LC is notably larger than the GL. The GL occupies the space of ∼2-5 LC neural tissue pores. The inset in the left panel shows the GL at the same scale as the LC. Scale bars: 250µm.

Genetically modified mouse models can allow excellent visualization of individual astrocyte structures in the GL. Yet, the same is not readily accomplished in large animal or human eyes. In the absence of endogenously expressed fluorescent reporters revealing cell morphology in situ, many studies utilize immunohistochemical means for astrocyte visualization. GFAP, a common astrocyte marker, is widely used to identify astrocytes.^40–45^ GFAP-labeling conducted in healthy and glaucomatous human donor eyes optic nerve heads^11^ revealed significant differences in the relative abundance and structure of GFAP in neural tissue pores. This suggests alteration in overall astrocyte organization, in situ, between health and glaucoma.

It is important to note that GFAP labeling reveals the structure of intermediate filaments in a fraction of astrocyte processes. It does not, however, reveal the full extent of astrocyte boundaries or morphology, as shown in the brain, retina, and mouse GL.^40–45^ In both the GL and LC, even with GFAP revealing only a portion of the cell, the density of astrocytes prevents understanding of where the boundaries of one astrocyte stop and its neighbor starts with light microscopy. Because of this, it is challenging to draw empirical conclusions about individual astrocyte morphologies or changes between health and glaucoma with confidence.

In summary, the morphology of individual LC astrocytes in the context of their native 3D environment is crucial to support retinal ganglion cell axons, but poorly understood. LC astrocytes are assumed to be represented by a prominent glaucoma research model, the mouse, but this has not been investigated in detail. To address this gap in understanding, we previously adapted a Multicolor DiOlistic labeling (MuDi) method, developed by Gan et al^46^, to visualize individual LC astrocyte morphologies, *in situ*. We demonstrated that MuDi-labeled cells with astrocyte morphologies were GFAP-positive, visualized across high order processes to their end-feet, and distinguished from neighboring astrocytes through multicolor labeling. This method worked across species with a LC, including goat, sheep, pig, and non-human primate. We additionally demonstrated that through confocal microscopy and 3D model generation, we were able to quantify morphological features of individual astrocytes at a cell-by-cell and branch-by-branch level.

In this work, we aimed to 1) characterize the morphology of individual astrocytes in the LC and 2) determine differences and similarities with those from the mouse GL. The scope of this work is focused on testing the hypothesis that there are significant morphological differences between astrocytes in the LC and in the GL. We demonstrate morphological differences between and similarities among astrocytes of the LC and GL that inform their potential for function (**Fig. 2**.)

**Figure 2:**
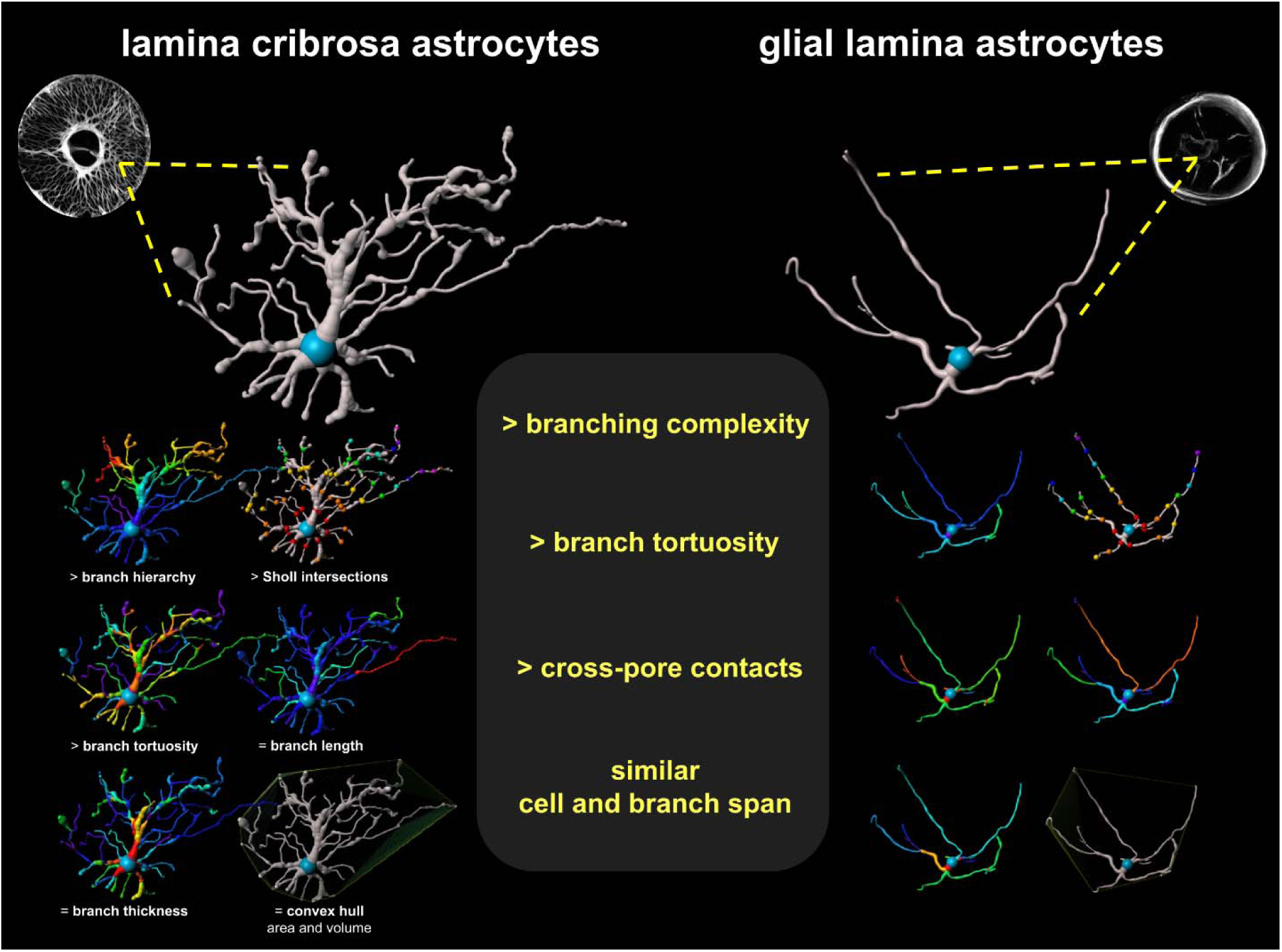
Astrocytes in the LC and GL had distinct and shared morphological features. LC astrocytes had greater branching complexity, branch tortuosity, and cross-pore contacts. LC and GL astrocytes had similar cell and branch spans.

## Methods

Twenty-two eyes with an LC were obtained from a local butcher within three hours of death (goat: N = 16, sheep: N = 4, pig: N = 2 eyes.) Methods for tissue preparation, imaging, and image processing were described in detail in our earlier work.^40^ We summarize these methods below.

### Tissue preparation

Optic nerve heads were isolated from globes, fixed in 4% PFA for 2-24 hours, embedded in low melting temperature agarose (Sigma, A9045), and sectioned into 150µm-thick slices with a vibratome (Leica, VT1200). Slices containing lamina cribrosa were stored at 4L in PBS for up to 48 hours. Individual astrocytes throughout tissues were stochastically labeled across their cell membranes with 7 distinct dye combinations via Multicolor DiOlistics^46^ (MuDi) as done previously.^40^ For MuDi labeling, gold microcarriers 1µm in diameter were coated with all combinations of 3 spectrally distinct dyes, DiI (Sigma-Aldrich 42364, 549/567 nm), DiO (Invitrogen D275, 484/501 nm), and DiD (Invitrogen D7757, 644/665 nm excitation/emission). Dye-coated microcarriers were loaded into Tefzel tubing (Bio-Rad, 1652441) and propelled into tissues at 150-200 PSI. Microcarriers which contacted individual astrocytes transferred dye to the cell membrane of those astrocytes, allowing for visualization.

### Imaging

Tissues were screened for extent of dye spread at 4-40x magnification via confocal microscopy (BX61; Olympus) before collection of volumetric scans quantitative analysis at 40x magnification. These images were collected at an XY image size of 1024×1024 or 800×800 pixels, and with a 0.5–2 μm Z step throughout the visible thickness of the samples. Volumetric images of healthy naive hGFAPpr-GFP mouse^47^ GL astrocytes collected via confocal microscopy were obtained from work conducted and published previously by Wang et al.^12^ 3D segmentations were created from images of 10 GL astrocytes from this earlier work.^12^

### Image processing

Images of LC astrocytes and mouse GL astrocytes were imported into Imaris software (Bitplane, version 9.9) for 3D visualization and generation of models. The Filaments tool and the “AutoPath-no loops” algorithm were used to automatically segment individual astrocytes. Each astrocyte was created with a single starting-point at its soma and seed-points along its branches. All Imaris-generated models were checked to make sure they represented image volume information faithfully. Aberrant branches were removed and missing branches were added manually. 40 3D models of goat LC astrocytes and 10 models of mouse GL astrocytes were generated for the core analysis of this study. Sholl analysis was conducted with the Filament Sholl Analysis XTension. Astrocyte convex hull area and volume were collected with the Filament Convex Hull function. Quantitative branch-by-branch level information about morphological features was extracted from the Statistics panel. These features were branch number, hierarchy, length, thickness, and straightness. Straightness is included as a % value, defined as the shortest distance between the two ends of the branch divided by the full path length of the branch. Branch tortuosity was calculated as 1-straightness, with values near 0% being straighter and values near 100% being more tortuous. Figures were made in part with BioRender.

### Feature selection

The morphological features selected for investigation were informed by features which undergo changes between health and glaucoma in mice. It has been previously demonstrated and/or suggested that the following astrocyte features differ between naive and glaucoma model mice: relationships with surrounding collagen^13^, Sholl area under the curve,^13^ convex polygon area,^13^ number of branches per astrocyte,^13^ branch hierarchy, tortuosity, thickness, length, and longitudinal processes.^12,13^ We address these differences in greater detail and what they indicate of astrocyte function in the Discussion.

### Statistical analysis

Statistical testing and plotting were completed in Python 3.1 through SciPy^48^, Matplotlib^49^, and seaborn^50^. Astrocyte centerpoints and branches with lengths of under 2µm, often erroneously produced at branch points, were filtered out of the dataset. Violin plots show data within the 95th percentile for each morphological feature evaluated. For each violin plot, the density distribution of data is shown in blue or red, the median is shown in white, the wider black line denotes the interquartile range, and thinner black lines denote the minimum and maximum values of the distribution, excluding outliers. For comparison of astrocyte features from the LC vs the GL, data were first analyzed for equality of variance and normality of distribution. Data with unequal variances and/or non-normal distributions were compared with a Kruskal–Wallis test. Data with normal distributions and equal variances were compared with an independent samples t-test.

## Results

We previously described MuDi labeling as a method to reveal LC astrocyte morphology *in situ*.^40^ High-resolution volumetric images of MuDi-labeled astrocytes allowed quantification of cell-by-cell and branch-by-branch morphometry. In this work, we leveraged these techniques to visualize and quantify astrocyte LC astrocyte morphologies. We characterized the morphology of individual LC astrocytes, in situ, and determined differences and similarities with astrocytes in the mouse GL. We show quantitative morphological differences between astrocytes of the LC and GL as well as features that were similar across the two.

### Model generation

We generated 40 astrocyte 3D models from images of MuDi-labeled^40,46^ goat LC (**Fig. 3**.) Astrocyte morphologies were diverse, similar to fibrous astrocyte morphologies in other areas of the central nervous system.^37^ Individual LC astrocytes occupied more space along the coronal plane (**Fig. 3A**) than the sagittal plane (**Fig. 3B**). Additionally, we generated 10 3D models from images of naive mouse GL astrocytes, previously investigated in Wang et al.^12^ We quantified morphologic features of 3561 branches from 40 LC astrocytes and 399 branches from 10 GL astrocytes. Models of 11 astrocytes from sheep LC and 5 from pig LC were collected as a preliminary test to investigate if there were any large differences in astrocyte morphology observed across LC astrocytes among different species.

**Figure 3:**
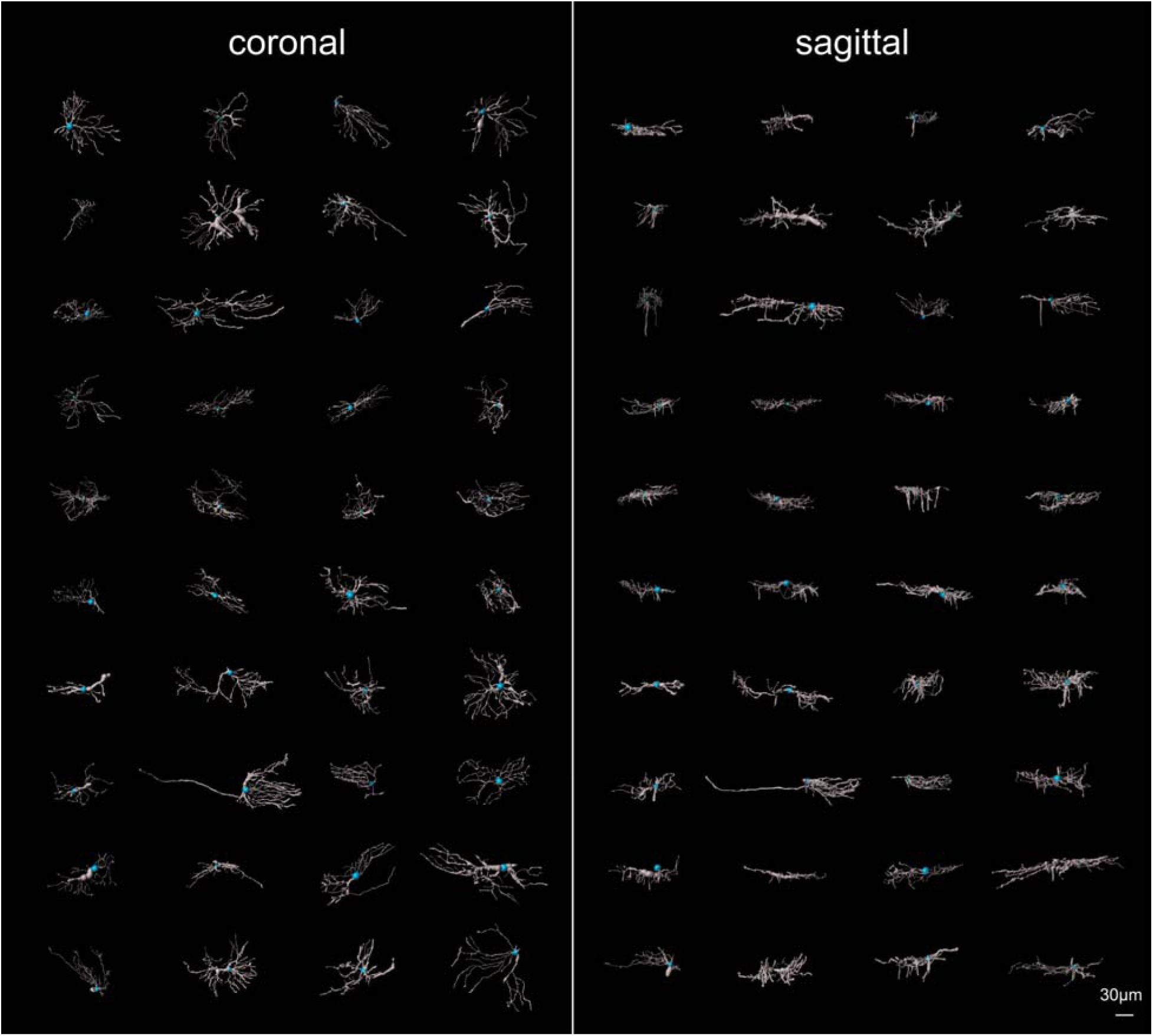
LC astrocyte morphologies are diverse. 40 image-derived 3D segmentations of goat LC astrocytes were created for morphometric analysis. Soma center-points are indicated in blue and astrocyte branches are indicated in gray. Left) Coronal view, right) sagittal view.

### Astrocyte relationships to collagen

One key anatomical difference between the LC and GL is the abundance and arrangement of collagen in the region. As the name connotes, the LC is a collagen-dense structure,^39,51–53^ composed of a network of collagenous beams which form pores occupied by neural tissue. In the GL, collagenous beams are absent; processes of individual astrocytes can span the width of the nerve and contact the collagenous canal perimeter.

In this work, we observed that the arrangement of astrocytes across the network of pores composing the LC is distinct from that of the GL. Astrocytes in the LC can be confined to individual neural tissue pores (**Fig. 4A**), similar to how GL astrocytes are confined to the width of the canal. Astrocytes in the LC can additionally span multiple pores (**Fig. 4B**), indicating potential for direct cell-cell communication and/or complex mechanosensation across pores. Given the anatomy of the GL, this arrangement of astrocytes is not possible in the mouse.

**Figure 4:**
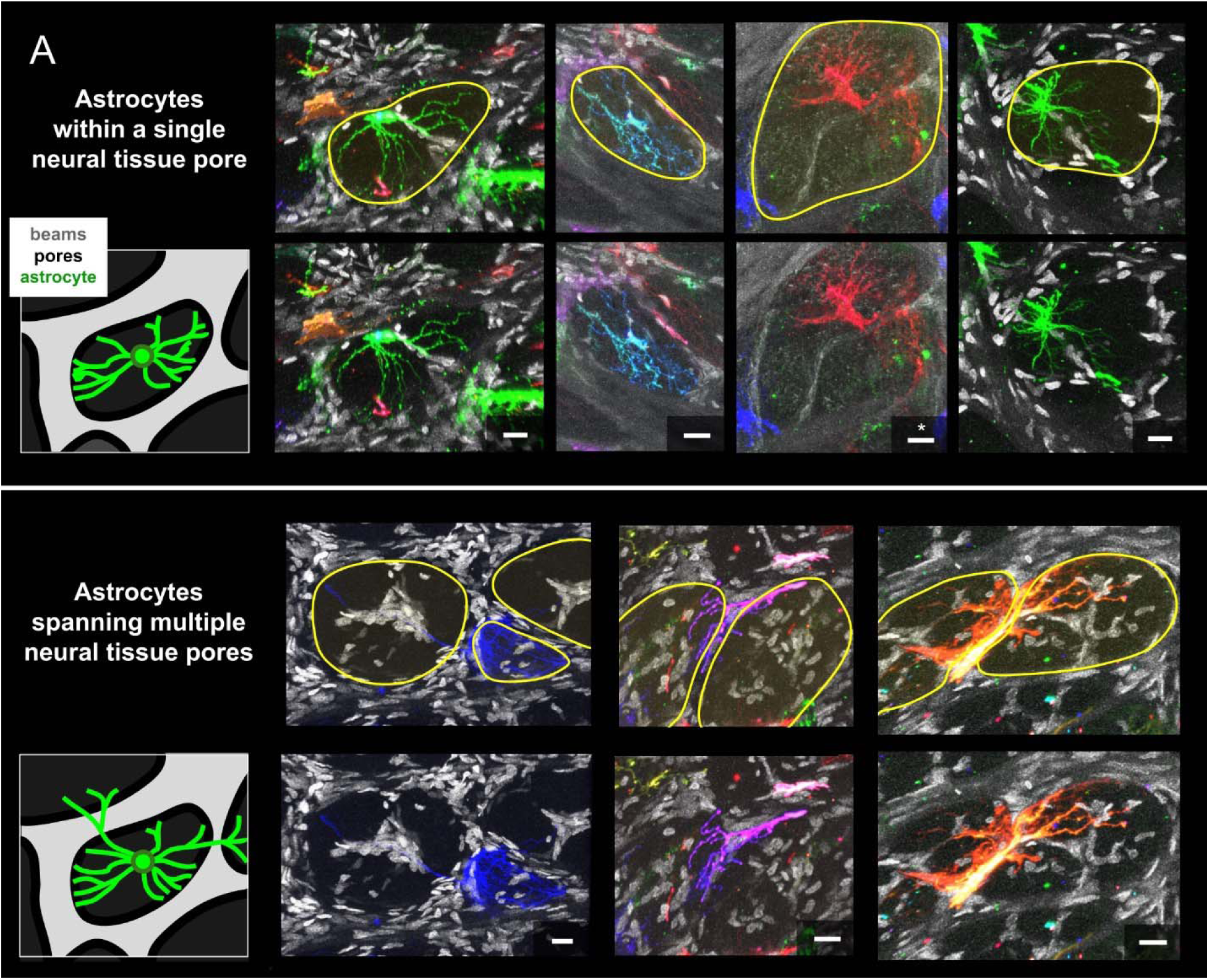
The arrangement of individual LC astrocytes within the network of LC collagenous beams. Astrocytes can be **A**) confined to a single neural tissue pore or **B**) span multiple pores. Top rows in **A** and **B** show neural tissue pore boundaries delineated in yellow. Bottom rows show astrocyte images without pore delineations. Astrocytes are shown in color. DAPI and collagenous beams are shown in grayscale. (Sample with *, second from the right in panel A, was not DAPI-labeled.) Dark regions indicate neural tissue pore areas without labeled astrocytes. Scale bars: 20µm.

### Sholl analysis

**Sholl analysis** was used to quantify branching complexity of individual astrocytes. 3D Sholl analysis quantifies the number of branch crossings along the edge of spheres with increasingly large radii, each centered at an astrocyte centerpoint. The more highly branched an astrocyte is at a given distance from its centerpoint, the higher number of crossings it will have. Sholl analysis (**Fig. 5**) revealed a significantly larger area under the curve for goat LC astrocytes compared to mouse GL astrocytes (LC = 709.4 ± 238.9 (mean ± SD), n = 40 cells, GL = 478.0 ± 238.9, **n =** 10 cells, p = 0.009, independent samples t-test), indicating higher astrocyte branching complexity in the LC. Mean number of total Sholl crossings for LC and GL astrocytes were 78.4 ± 24.1 and 52.1 ± 26.8, respectively.

**Figure 5:**
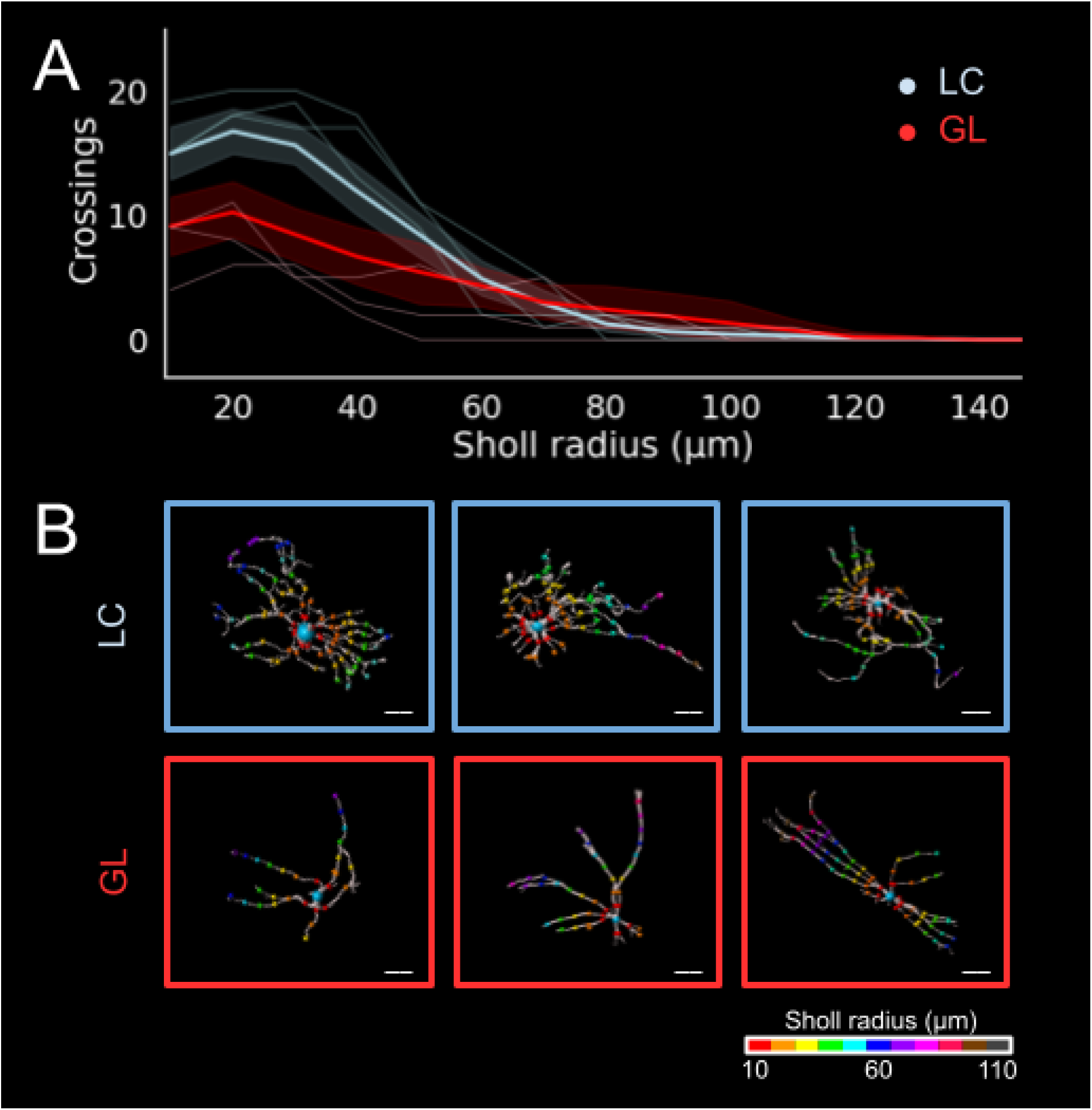
Sholl analysis reveals higher branching complexity in LC astrocytes than in GL astrocytes. Branching complexity was measured via 3D Sholl analysis. Area under the Sholl curve was compared between LC and GL astrocytes. **A**) Bold lines: mean crossings at each Sholl sphere radius, shaded areas: standard deviation. Thin lines show traces from example individual cells, shown below in **B**. Scale bars: 20µm.

Maximum Sholl span signifies the length that astrocytes can span from centerpoint to distal tip. Despite substantial differences in the anatomy and size of the LC and the GL, maximum Sholl span of LC and GL astrocytes was not significantly different (maximum Sholl span per cell; LC: 77.9 ± 26.6µm, GL: 86.0 ± 26.3, p = 0.352, independent samples t-test.)

### Astrocyte branch features

**Number of branches** per astrocyte (**Fig. 6**) was significantly higher in the LC (86.4 ± 25.3 branches) than in the GL (39.3 ± 18.5 branches, p < 0.001, independent samples t-test) by 119.8%. **Branch hierarchy** of astrocytes (**Fig. 7**) was significantly higher in the goat LC (6.6 ± 3.5µm) than the mouse GL (4.2 ± 2.0µm, p < 0.001, Kruskal-Wallace test) by 2.4, 57.14% higher). Maximum branch hierarchy was 22 for LC astrocytes and 12 for mouse GL astrocytes. **Branch tortuosity** of LC astrocytes (**Fig. 8**, 12.3 ± 9.2%) was significantly higher than in the GL (7.3 ± 6.7%, p < 0.001, Kruskal-Wallace test), with LC tortuosity 68.5% higher GL tortuosity. Astrocyte **branch length** (**Fig. 9**) was not significantly different between those from the LC (13.7 ± 11.5µm) and the GL (17.0 ± 17.2µm, p = 0.10, Kruskal-Wallace test.) Maximum LC astrocyte branch length was 179.6µm. Astrocyte **branch diameter** (**Fig. 10**) was not significantly different between the LC (1.8 ± 1.1µm) and the mouse GL (1.8 ±1.0µm, p = 0.64, Kruskal-Wallace test).

**Figure 6:**
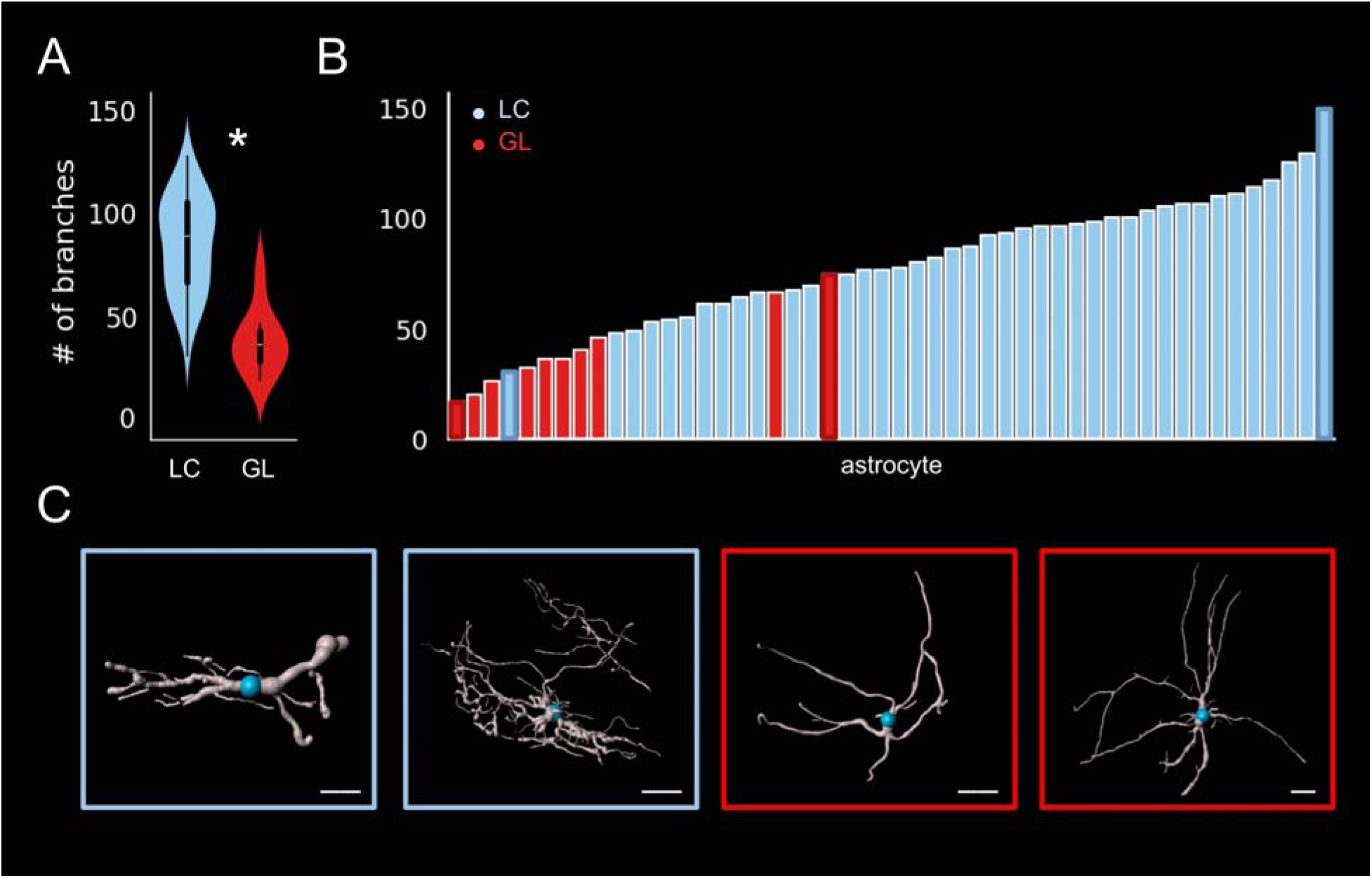
LC astrocytes have more branches than GL astrocytes. **A**) Distribution of number of branches per astrocyte shown in a violin plot, * connotes a significant difference between the LC and GL, p < 0.05. **B**) Number of branches per astrocyte, sorted from low to high. **C**) Example LC (blue) and GL (red) astrocytes with the lowest (left) and highest (right) number of branches. Scale bars: 20µm.

**Figure 7:**
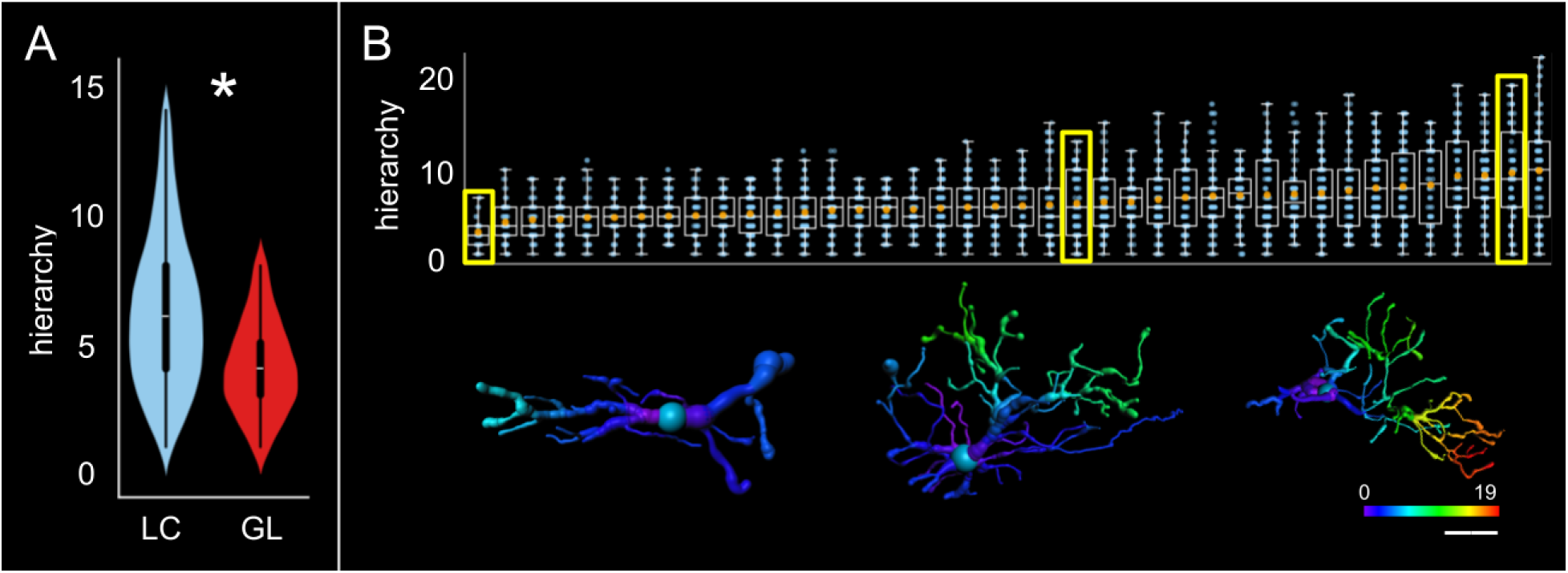
Branch hierarchy is deeper in LC astrocytes than GL astrocytes. **A)** Branch hierarchy was significantly deeper in LC astrocytes than GL astrocytes (p < 0.001.) **B)** Branch hierarchy of LC astrocytes by cell. Each box plot represents an individual astrocyte, blue points each represent a branch, and orange points represent mean branch hierarchy per astrocyte. Note that measurements of hierarchy are integers, which results in clusters of points in the box plots. The points in these clusters do not look identical because the plotting technique uses horizontal jitter and partial transparency to help more clearly display multiple points corresponding with branches of the same hierarchy. Example astrocytes with low, near-middle, and high mean branch hierarchy shown below. Their respective box plots are highlighted in yellow. Scale bar: 20µm.

**Figure 8:**
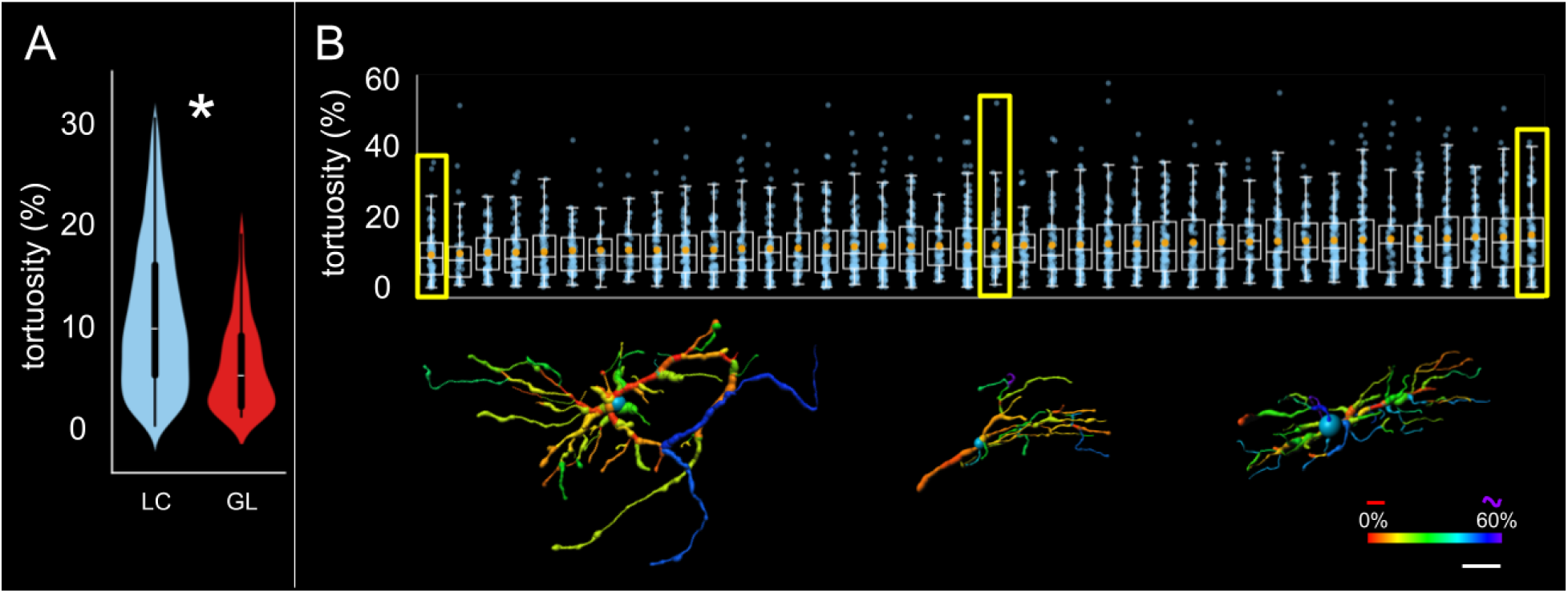
Branch tortuosity is higher in LC astrocytes than GL astrocytes. **A)** Branch tortuosity was significantly higher in LC astrocytes than GL astrocytes (p < 0.001.) **B**) Branch tortuosity of LC astrocytes by cell. Each box plot represents an individual astrocyte, blue points each represent a branch, and orange points represent mean branch hierarchy per astrocyte. Example astrocytes with low, near-middle, and high mean branch tortuosity shown below and their respective box plots highlighted in yellow. Scale bar: 20µm.

**Figure 9:**
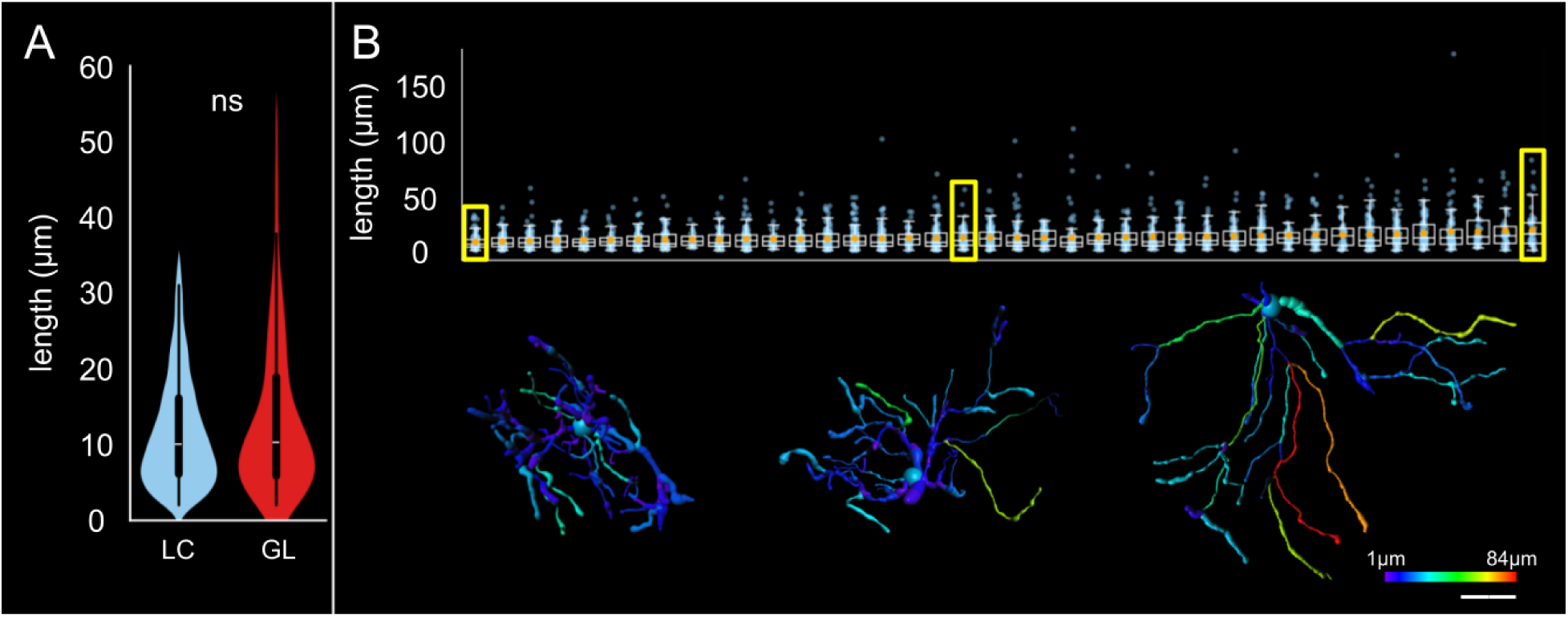
Branch length was not significantly different between LC and GL astrocytes. **A)** Branch length of LC astrocytes was not significantly different between LC and GL astrocytes (p = 0.10.) **B**) Branch length of LC astrocytes by cell. Each box plot represents an individual astrocyte, blue points each represent a branch, and orange points represent mean branch hierarchy per astrocyte. Example astrocytes with low, near-middle, and high mean branch length shown below and their respective box plots highlighted in yellow. Scale bar: 20µm.

**Figure 10:**
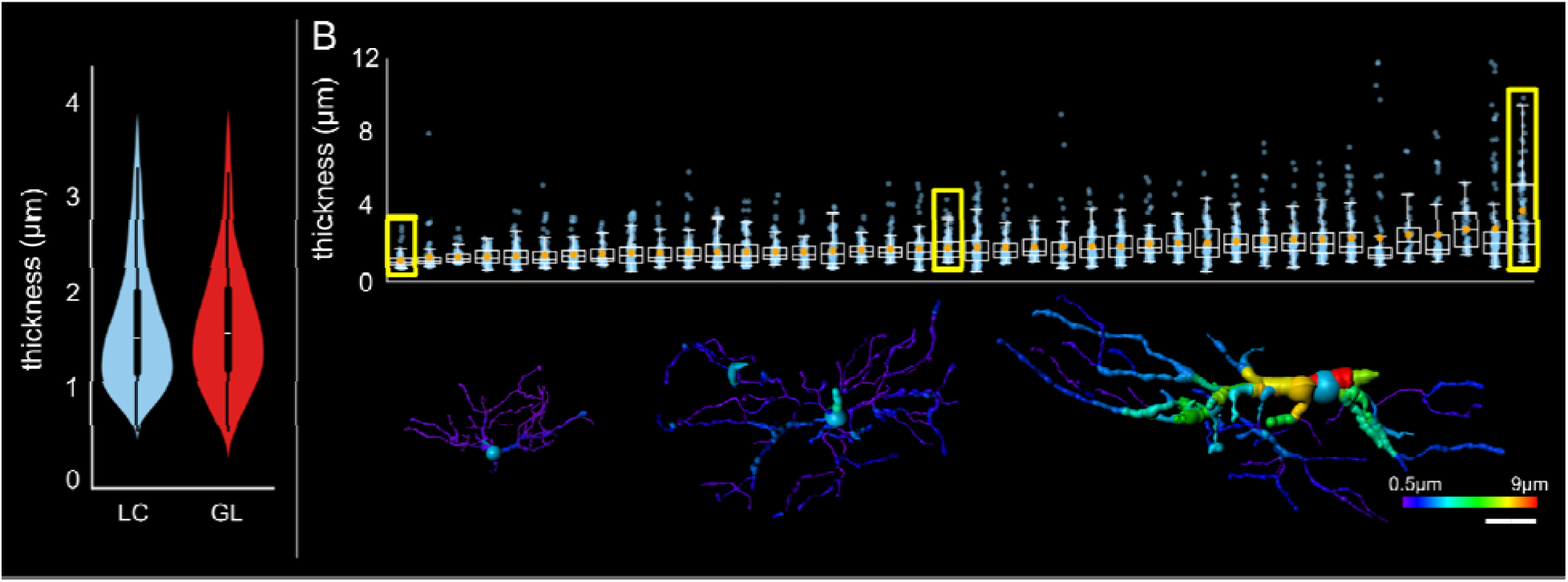
Branch thickness of LC astrocytes was not significantly different between LC and GL astrocytes. **A**) Branch length of LC astrocytes was not significantly different between LC and GL astrocytes. (p > 0.05.) **B**) Branch thickness of LC astrocytes by cell. Each box plot represents an individual astrocyte, blue points each represent a branch, and orange points represent mean branch hierarchy per astrocyte. Example astrocytes with low, near-middle, and high mean branch thickness shown below and their respective box plots highlighted in yellow. Scale bar: 20µm.

### Astrocyte spatial territory / span

Astrocyte convex hulls were created to gain a quantitative understanding of cellular spatial territories. Astrocytes of the LC and GL were found to occupy similar extents of spatial territory (**Fig. 11**.) Convex hull area (LC: 1.8×10^4^ ± 7.0×10^3^ µm^2^, GL: 2.0×10^4^ ± 1.7×10^4^ µm^2^, **Fig. 11A**) was not significantly different among LC and GL astrocytes. Convex hull volume (LC: 1.3×10^5^ ± 7.8×10^4^, GL: 1.4×10^5^ ± 1.8×10^5^, **Fig. 11B**) was also not significantly different among LC and GL astrocytes. Convex hulls tended to occupy more space along the coronal plane than the sagittal one (**Fig. 11C**), consistent with astrocyte models shown **Fig. 3**. Additionally, astrocytes in the LC were found to occupy shared spatial territories (**Fig. 12**), like those in the GL.^38^ This is in contrast with astrocytes in other regions of the central nervous system, which demonstrate a tiled arrangement with limited spatial overlap.^41,43,54^

**Figure 11:**
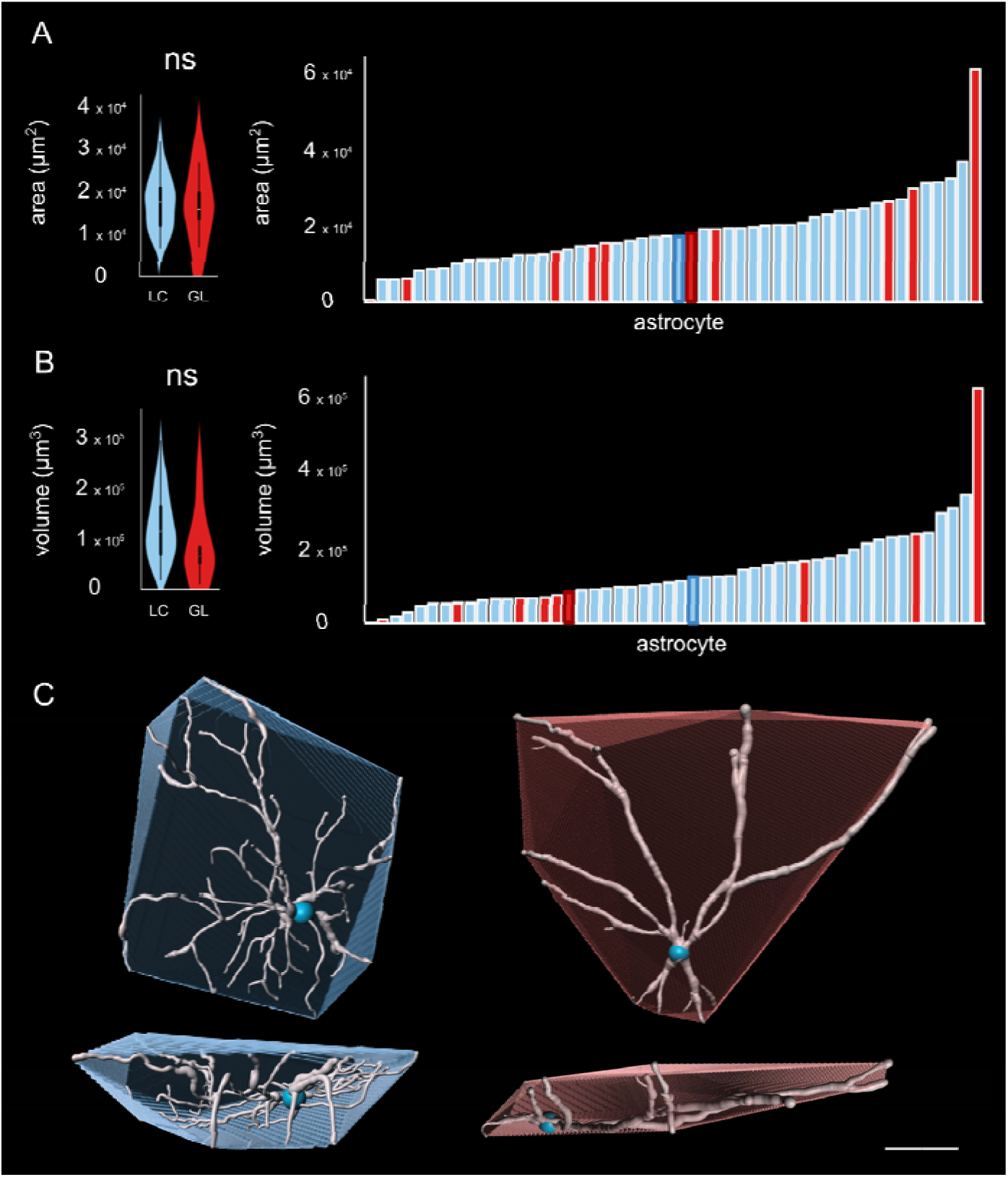
Astrocytes in the LC and GL occupy similar degrees of spatial territory. Astrocyte spatial territories were quantified as convex hull area and volume. Convex hull **A**) area and **B**) volume were not significantly different between the LC and GL (p = 0.98 and 0.21, respectively). Violin plots show aggregate data distribution and bar charts show cell-by-cell distribution. Bars bolded in **A** indicate representative LC and GL astrocyte convex hulls shown in **C.** LC: blue, GL: red. Scale bar: 20µm.

**Figure 12:**
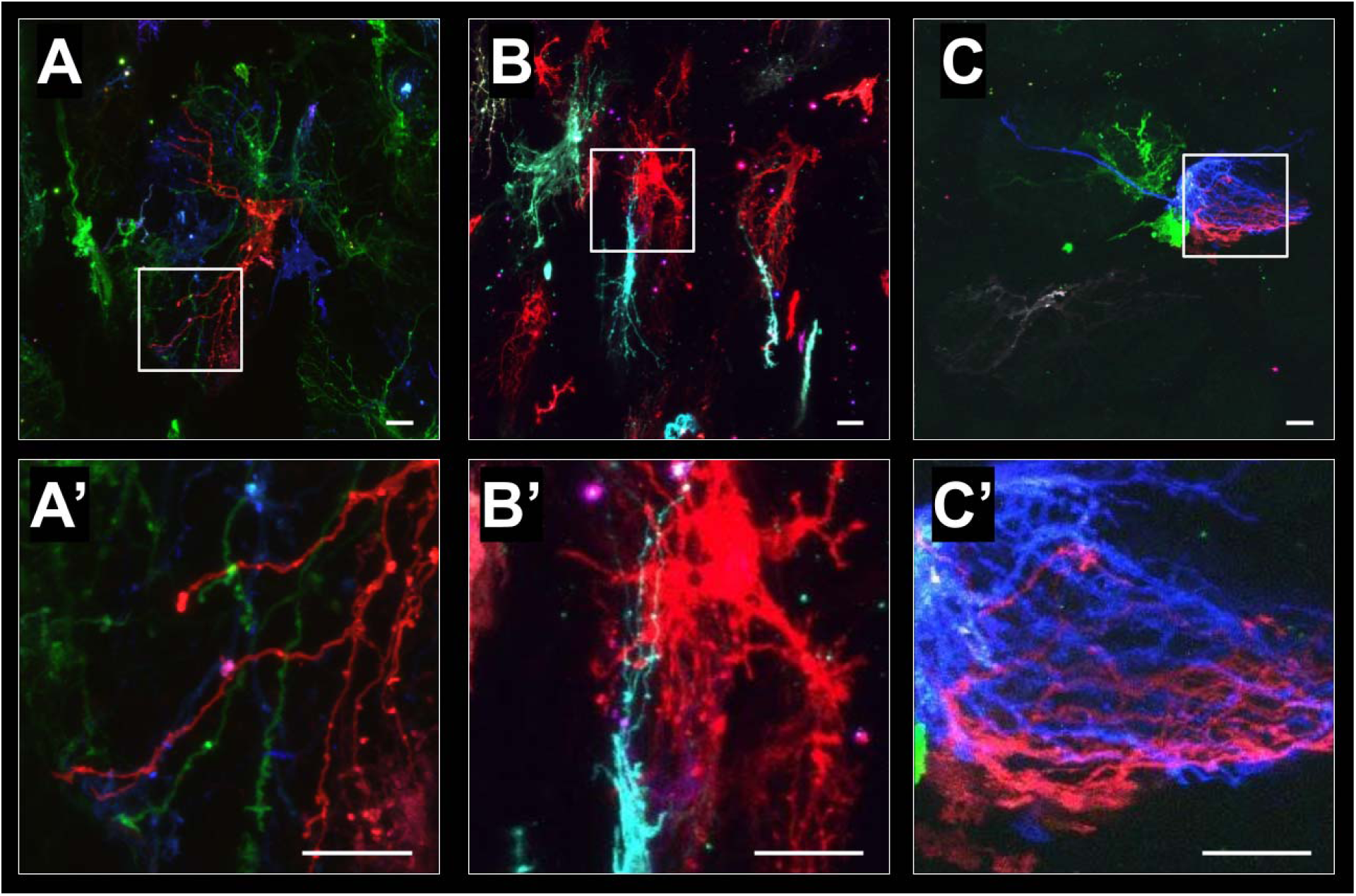
LC astrocytes demonstrate shared spatial territories. In contrast with other central nervous system tissues, astrocytes in the LC do not demonstrate distinct spatial domains or tiling. Overlapping astrocyte spatial territories has been demonstrated previously in the mouse GL. Example LC astrocytes with overlapping spatial territories are shown in **A**-**C**. Detail of regions where astrocytes overlap is shown in **A’**-**C’**. Scale bars: 20µm.

### Longitudinal processes

Although most GL astrocyte branches are oriented along the coronal plane, longitudinally-oriented branches have been observed.^12,13^ In LC astrocytes, where most astrocytes branches are also primarily oriented along the coronal plane (**Fig. 3**), we observed astrocytes with similar longitudinally-oriented branches. Example LC astrocytes with longitudinally-oriented branches color-coded by their position along the anterior-posterior axis are shown in **Fig. 13**.

**Figure 13:**
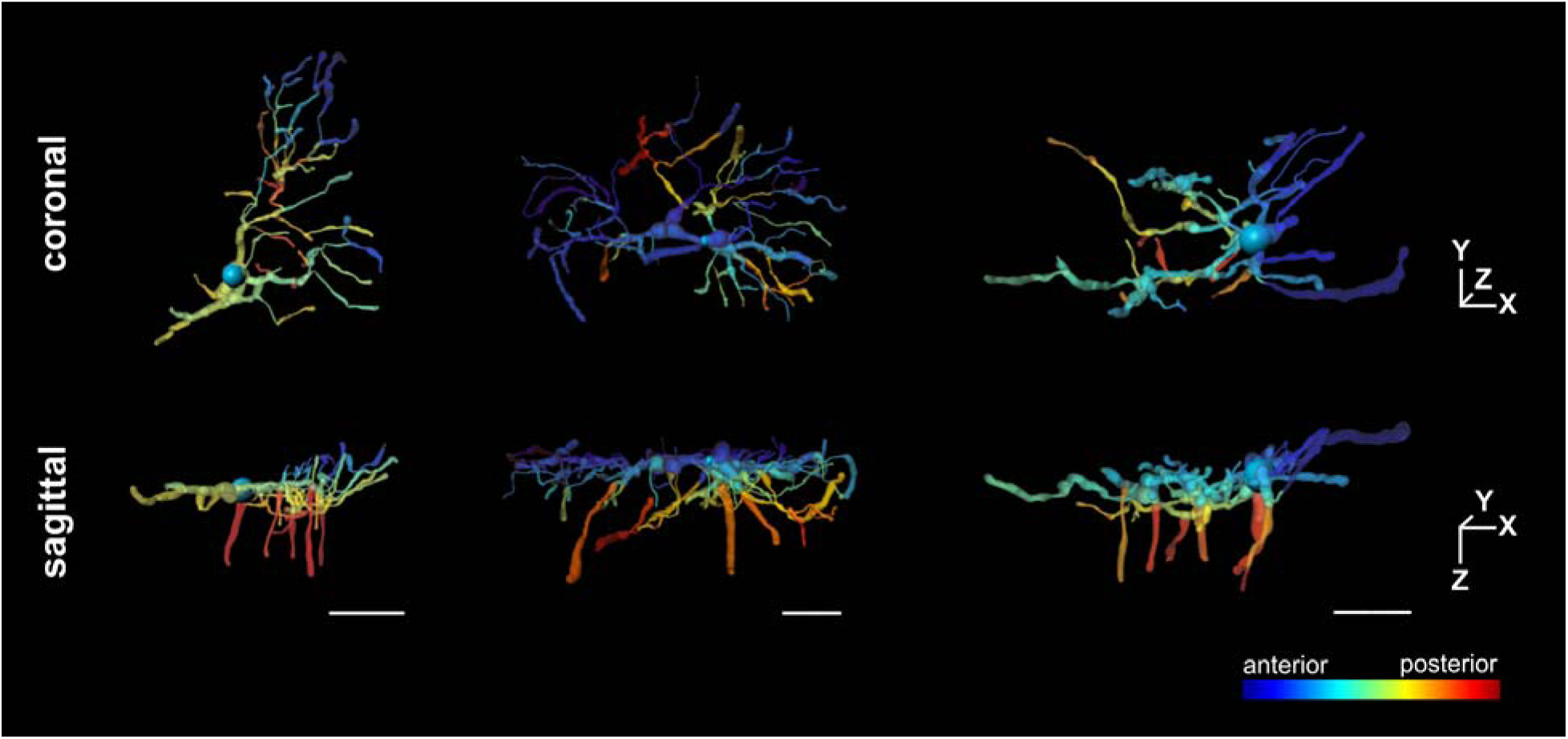
Longitudinal processes of LC astrocytes. Example LC astrocytes with processes oriented primarily along the anterior-posterior direction. Color scale denotes astrocyte branch location along the anterior-posterior axis. Scale bars: 20µm.

### Cross-species LC comparisons

Our preliminary data on astrocyte branch features in other species with a LC suggests that goat LC astrocytes represent astrocytes of other species with a LC well (**Fig. 14**.) These findings suggest that LC astrocyte morphologies may generally be more similar to each other than they are to GL astrocyte morphologies. This must be investigated further with a dataset of more astrocytes across species with a LC.

**Figure 14:**
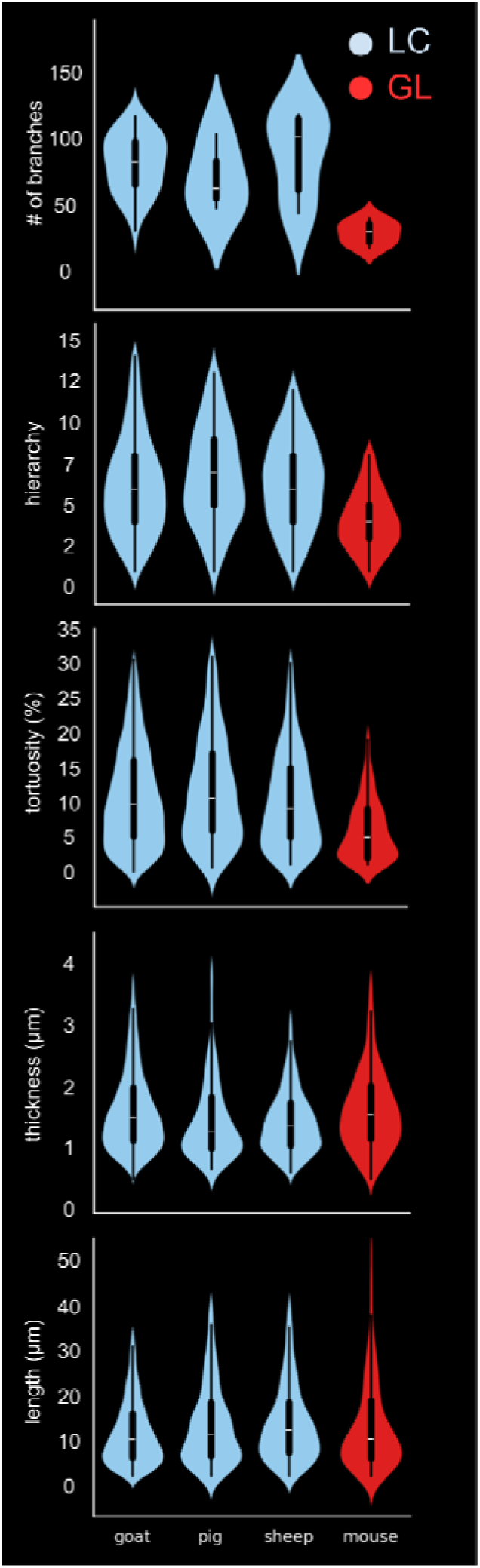
Differences and similarities between the GL and LC branch features appear consistent across species. Branch features of all astrocytes quantified from species with a LC (blue) and from the GL (red) demonstrate similar relationships of goat vs. mouse with a more general pattern of LC vs GL. Mean number of branches per astrocyte, branch hierarchy, and branch tortuosity were higher in all species with a LC compared to mouse. Mean branch thickness and length were relatively similar among species with a LC and with mouse.

## Discussion

In this work, we quantified morphological features of astrocytes in the LC to provide an in depth characterization of the morphology of these cells, in situ. Additionally, we compared the morphology of LC astrocytes with the morphology of astrocytes in an analogous structure, the GL from the mouse, a common glaucoma research model. This has revealed key findings related to differences between LC and GL astrocytes, and similarities between them. Astrocytes of the LC and GL differed substantially in **1)** their spatial relationships to surrounding collagen, **2)** branching complexity, and **3)** branch tortuosity. Regarding similarities, LC and GL astrocytes had **4)** similar cell and branch span, despite differences in the size of their anatomical environments.

### Differences between LC and GL astrocyte structures

#### Spatial relationships with collagen

The anatomies of the GL and the LC are distinct, particularly in relation to their size and organization of collagen. The mouse GL spans ∼200-300µm along the coronal direction.^38^ GL astrocytes typically span at least half the diameter of the canal and have numerous contacts with its collagenous perimeter.^38^ The GL lacks collagenous beams^55^ which divide neural tissue into pores, characteristic of the LC.

The LC is present in the eyes of large mammals, including humans. It spans 1.3-2.2mm^56,57^ along the coronal direction in humans and is generally slightly larger in ungulates.^51,52,58^ Average pore diameter in the human and non-human primate LC is ∼25µm and again slightly larger in ungulates.^33,34,58–61^ Limited knowledge is available about the arrangement of individual astrocytes in the LC or how this compares to the GL. Other work has characterized the abundance and distribution of nuclei and GFAP in serial sections of human donor eyes.^11^ This has provided an understanding of how many astrocytes exist in the LC and the extent of their GFAP-positive coverage. Given the abundance of GFAP in the LC, it is not possible to distinguish individual astrocytes. Additionally, GFAP does not reveal the full extent of astrocyte morphology.^40–45^

In our work, we observed individual LC astrocytes with variable spatial relationships to LC collagenous beams. Some astrocytes were confined to individual neural tissue pores, with many branches contacting pore perimeters. The arrangement of these astrocytes within individual neural tissue pores is similar to the arrangement of astrocytes within the entirety of the mouse GL. Additionally, we observed LC astrocytes spanning across neural tissue pores. These LC astrocyte spatial relationships to collagen are unique from those in the GL.

The ability of LC astrocytes to span multiple pores has interesting implications for local mechanosensation and cell signaling. Because the arrangement of collagen influences the biomechanics of the LC,^58,62,63^ the experiences of astrocytes at healthy and glaucomatous IOPs may have important differences between the GL and LC. Future work must be done to determine how LC astrocytes with different spatial relationships to collagenous beams respond to different IOPs. As glaucomatous damage to RGC axons can be focal^64^ and astrocytes can play a role in this damage^65^, future work can additionally be directed at understanding LC astrocyte signaling within pores and across them.

#### Branching complexity

Overall, each metric related to branching complexity collected was found to be higher in the LC compared to the GL. Sholl analysis indicated a significantly higher area under the curve and a higher number of Sholl crossings per cell in the LC than in the GL. The number of branches per astrocyte was higher in the LC than the GL. Branch hierarchy was deeper in the LC compared to the GL.

Higher astrocyte branching complexity has been documented in central nervous system tissues of larger mammals, including non-human primates and humans, compared to mouse.^37^ These structural differences were coincident with functional differences. For example, the rate of astrocyte calcium wave transmission was significantly faster in humans than in mice.^37^ Human astrocytes were more susceptible to oxidative stress than mouse astrocytes.^66^ As both calcium signaling and oxidative stress are implicated in glaucoma, it can be valuable to look further into how mouse GL astrocytes may differ functionally from those in larger mammals, including humans.

Branch complexity has been observed to decrease in the mouse GL under hypertensive conditions.^13^ Decreased branch complexity provides fewer opportunities for astrocyte contacts with surrounding collagen, blood vessels, axons, and other astrocytes. This is likely to impact their capacity for sensing and signaling, contributing to glaucomatous dysfunction. The extent to which astrocyte branching simplification takes place in glaucoma in larger eyes is not known. It’s possible that increased branch complexity of LC astrocytes provides a degree of protective redundancy. Alternatively, it may indicate distinct physiological requirements for healthy function.

#### Branch tortuosity

Images from studies of the GL suggest that under hypertensive conditions, GL astrocyte branches become straighter.^13^ Additionally, these astrocytes express mechanosensitive channels^10^, indicating that they have the capacity for mechanosensation. In both the GL and LC, moderate IOPs typically do not cause RGC injury. However, large IOP increases can lead to astrocyte activation, contributing to RGC axon damage.^1,2^ Astrocyte branch tortuosity may provide mechanical slack, influencing the effects of IOP-induced stretch on astrocyte mechanosensation. Astrocyte branches with lower tortuosity may be more susceptible to IOP-induced stretch beyond their degree of slack, and therefore potentially more susceptible to mechanical damage (**Fig. 15**).^67–69^ In cases of glaucoma with normal or only moderately elevated IOP, low astrocyte branch tortuosity may be an indicator of individual sensitivity to IOP. Astrocytes in the LC had significantly higher branch tortuosity than GL astrocytes. This difference may have important mechanical implications in the context of health and glaucoma.

**Figure 15:**
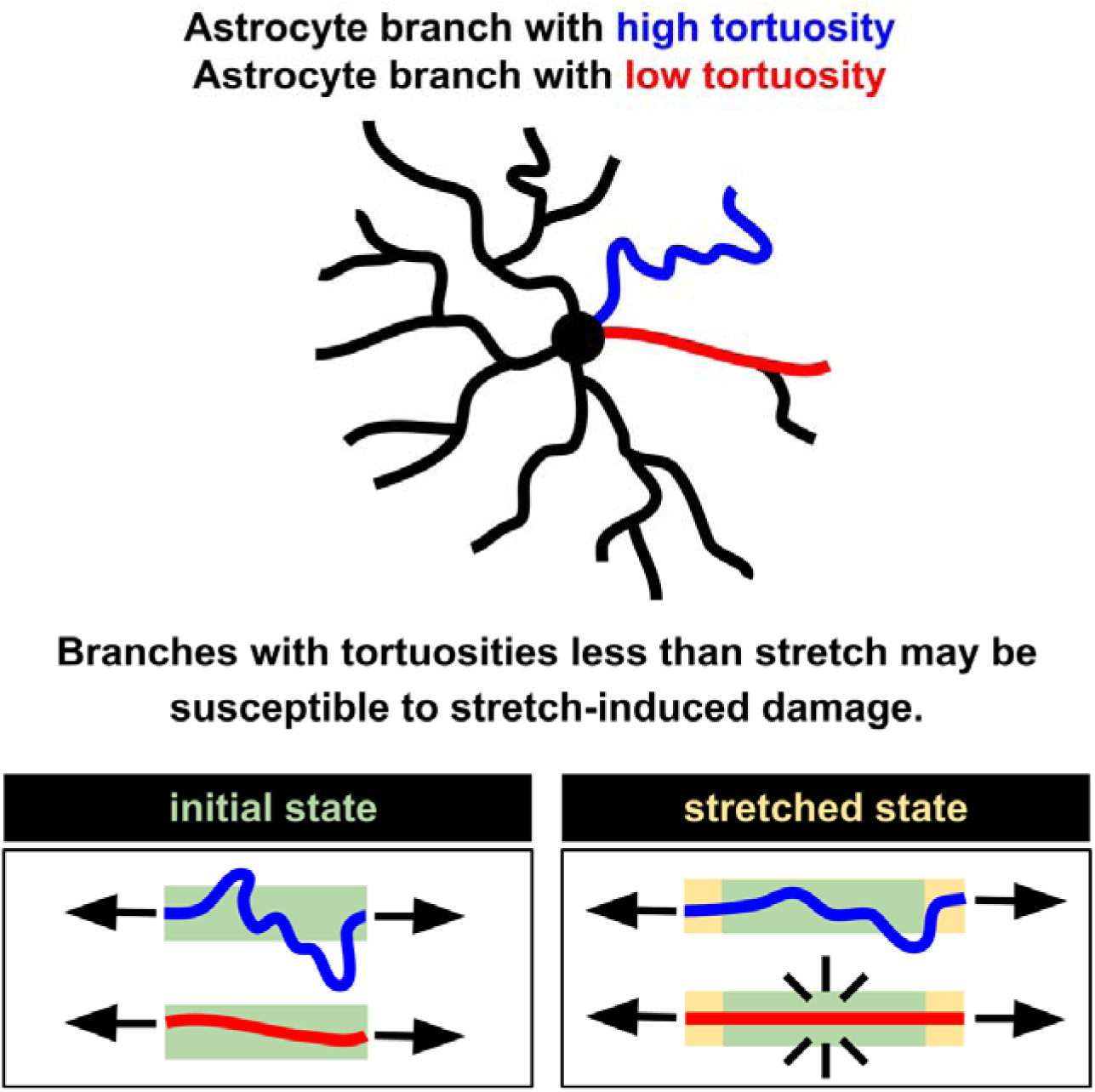
Astrocyte branch tortuosity as a mechanism to withstand IOP-induced insult. Astrocyte branches with slack to withstand IOP-induced stretch may be mechanically protected in comparison to astrocyte branches that do not have the degree of slack to withstand IOP-induced stretch. Example astrocyte branches with high and low tortuosity are shown in blue and red, respectively.

Collagen is a key load-bearing component of the LC. Collagen fibers have a natural degree of tortuosity or “crimp” that influences their biomechanics. As IOP increases, crimped LC collagen has been shown to straighten and become stiffer.^62,70,71^ With LC astrocyte branch tortuosity from this work computed as arc length / chord length, as in other work focused on LC collagen crimp,^58^ mean astrocyte branch tortuosity was 1.155 ± 0.146. In work using sheep eyes fixed at 5mmHg IOP, mean LC collagen crimp tortuosity was 1.017 ± 0.028 for thin beams and 1.025 ± 0.037 for thick beams.^58^ This LC collagen tortuosity is lower than LC astrocyte branch tortuosity. This may suggest that LC collagen will generally straighten and stiffen before astrocyte branches are stretched beyond their degree of slack. A mismatch between local LC collagen tortuosity and local LC astrocyte branch tortuosity could indicate predisposition to mechanical insult.

It must be noted that all LC and GL astrocytes analyzed in this work were fixed and imaged at 0mmHg IOP. Branch tortuosity at higher IOPs is expected to be lower than at this unpressurized baseline. In future work, it will be valuable to compare IOP-induced changes in LC collagen tortuosity directly with changes in astrocyte branch tortuosity in the same tissues.

### Similarities between LC and GL astrocyte structures

#### Cell and branch span

Despite anatomical environments that are substantially different in size and collagen composition, LC and GL astrocytes demonstrated similar cell and branch spans. Astrocyte cell span (indicated by maximum Sholl sphere crossing radius, convex hull area, and convex hull volume) and branch span (indicated by branch thickness and length) were not significantly different between the LC and GL.

Why might this be unexpected? The coronal diameter of individual neural tissue pores in the LC is generally smaller than that of the GL.^33,34,38,58–61^ If astrocytes within LC neural tissue pores were organized similarly to those within a single mouse GL, with astrocyte endfeet touching pore edges in the LC and canal edges in the GL, LC astrocytes would span less distance than GL astrocytes. In other central nervous system tissues, mouse fibrous astrocytes have been shown to span less area than fibrous astrocytes from larger mammals, such as humans and non-human primates.^37^ We observed that LC astrocytes have similar cell spans to GL astrocytes. This is in part due to LC astrocytes often spanning multiple LC pores. Individual branches of LC and GL astrocytes had similar spans as well. No significant differences were found between LC and GL astrocytes in branch thickness or length.

From studies conducted in a mouse model of glaucoma, elevated IOP has been shown to affect astrocyte cell span and branch span. With elevated IOP, astrocyte cell span decreased, branch thickness increased, and representative images suggest a substantial decrease in branch length.^13^ Future work must be done to determine whether the IOP-induced astrocyte changes observed in the GL are similar or different in the LC.

Astrocytes and most of their branches largely spanned along the coronal axis. Interestingly, we observed astrocyte longitudinal processes, similar to those documented in the mouse GL.^12,13^ The function of these longitudinal processes is not well understood. Their presence in astrocytes of both the GL and LC may implicate similar roles in both structures.

#### Strengths of this work

The GL is a strong and accessible model for many aspects of glaucoma research. Mouse lines with genetically-encoded fluorescent reporters, such as those used to investigate GL astrocytes, are excellent tools for understanding the structure and organization of cells in situ. However, the GL cannot be assumed to fully represent the LC. Methodological limitations have prevented the study of individual astrocyte morphologies in the LC. We demonstrated MuDi as a method to allow visualization and quantitative morphological evaluation of individual astrocytes in the LC.^40^ Leveraging this approach to focus on astrocytes in the LC, which are poorly understood compared to those in the GL, is a strength of this work.

Reports of astrocyte morphology in health and disease outside of mouse models often rely on immunohistochemical labeling of GFAP. The high density of astrocytes in the LC prevents distinguishing individual astrocytes through GFAP labeling. Additionally, GFAP does not reveal the complete boundaries of the cell, providing limited information about astrocyte morphology. GFAP expression and organization is influenced by stressors, including elevated IOP.^72–74^ However, it’s not known whether these differences in GFAP organization represent changes in cell morphology well. MuDi utilizes cell membrane labeling to reveal the full exterior boundaries of the cell. We provide quantitative morphological information that doesn’t rely on incomplete and variable marking of astrocyte boundaries by GFAP.

#### Limitations

Important limitations of this work must be noted. The DiOlistic labeling used in this work relies on stochastic transfer of cell membrane dyes to a subset of cells. This, along with a multicolor labeling approach, allowed us to distinguish individual astrocytes from their neighbors for analysis of individual cell morphologies. This leaves many astrocytes unlabeled. We were therefore not able to visualize all astrocytes across the LC. Additionally, we used coronal vibratome sections for labeling. Because astrocytes that are larger and/or more coronally-oriented occupy more space along the plane of labeling, DiOlistically-labeled astrocytes may skew slightly larger and more coronally oriented than the overall population of astrocytes in the LC.

Our preliminary evidence on astrocyte morphology across species with a LC suggest LC-specific patterns. For example, mean number of branches, branch hierarchy, and branch tortuosity were higher in all species with a LC compared to the GL. Branch thickness and length were similar across all species. However, findings related to LC astrocytes presented here are primarily focused on data from the goat LC. As MuDi allows visualization of individual LC astrocytes in a species-agnostic manner,^40^ future work can be focused on non-human primate and human astrocytes. It will be valuable to determine if there are human-specific LC astrocyte features that are not captured by studying other species with a LC.

This work investigated the morphology of LC without the introduction of any glaucomatous stressors. This information about the structure and organization of LC astrocytes under normal conditions has provided important insights into their potential for function in health and disease and how this may differ between the LC and GL. Understanding the differences between healthy and glaucomatous LC astrocytes is outside the scope of this study. Future studies will be designed to determine the morphologic changes LC astrocytes undergo under glaucomatous conditions.

### Conclusions

In summary, we utilized MuDi, confocal microscopy, and 3D morphometric analysis to characterize individual astrocyte morphologies in the LC, in situ. We compared morphological features of LC and GL astrocytes to identify similarities and differences. Morphological features analyzed indicate capacity for function in health and glaucoma. Astrocytes in the LC had different spatial relationships with collagen, higher branching complexity, and higher branch tortuosity than GL astrocytes. Astrocytes in the LC and GL had similar cell and branch spans despite substantial differences in their anatomic environments. Further work is needed to better understand the potentially distinct roles of LC astrocytes in the initiation and progression of glaucoma.

## References

1. Stowell C, Burgoyne CF, Tamm ER, Ethier CR, Lasker/IRRF Initiative on Astrocytes and Glaucomatous Neurodegeneration Participants. Biomechanical aspects of axonal damage in glaucoma: A brief review. Exp Eye Res. 2017;157:13–19.

2. Tamm ER, Ethier CR, Lasker/IRRF Initiative on Astrocytes and Glaucomatous Neurodegeneration Participants. Biological aspects of axonal damage in glaucoma: A brief review. Exp Eye Res. 2017;157:5–12.

3. Verisokin AY, Verveyko DV, Postnov DE, Brazhe AR. Modeling of Astrocyte Networks: Toward Realistic Topology and Dynamics. Front Cell Neurosci. 2021;15:645068.

4. Boal AM, Risner ML, Cooper ML, Wareham LK, Calkins DJ. Astrocyte Networks as Therapeutic Targets in Glaucomatous Neurodegeneration. Cells. 2021;10(6). doi:10.3390/cells10061368

5. Fields RD, Woo DH, Basser PJ. Glial Regulation of the Neuronal Connectome through Local and Long-Distant Communication. Neuron. 2015;86(2):374–386.

6. Petzold GC, Murthy VN. Role of astrocytes in neurovascular coupling. Neuron. 2011;71(5):782–797.

7. Stackhouse TL, Mishra A. Neurovascular Coupling in Development and Disease: Focus on Astrocytes. Front Cell Dev Biol. 2021;9:702832.

8. Wan Y, Wang H, Fan X, et al. Mechanosensitive channel Piezo1 is an essential regulator in cell cycle progression of optic nerve head astrocytes. Glia. 2023;71(5):1233–1246.

9. Li Z, Peng F, Liu Z, Li S, Li L, Qian X. Mechanobiological responses of astrocytes in optic nerve head due to biaxial stretch. BMC Ophthalmol. 2022;22(1):368.

10. Choi HJ, Sun D, Jakobs TC. Astrocytes in the optic nerve head express putative mechanosensitive channels. Mol Vis. 2015;21:749–766.

11. Guan C, Pease ME, Quillen S, et al. Quantitative Microstructural Analysis of Cellular and Tissue Remodeling in Human Glaucoma Optic Nerve Head. Invest Ophthalmol Vis Sci. 2022;63(11):18.

12. Wang R, Seifert P, Jakobs TC. Astrocytes in the Optic Nerve Head of Glaucomatous Mice Display a Characteristic Reactive Phenotype. Invest Ophthalmol Vis Sci. 2017;58(2):924–932.

13. Lye-Barthel M, Sun D, Jakobs TC. Morphology of astrocytes in a glaucomatous optic nerve. Invest Ophthalmol Vis Sci. 2013;54(2):909–917.

14. Sun D, Moore S, Jakobs TC. Optic nerve astrocyte reactivity protects function in experimental glaucoma and other nerve injuries. J Exp Med. 2017;214(5):1411–1430.

15. Quillen S, Schaub J, Quigley H, Pease M, Korneva A, Kimball E. Astrocyte responses to experimental glaucoma in mouse optic nerve head. PLoS One. 2020;15(8):e0238104.

16. Almasieh M, Wilson AM, Morquette B, Cueva Vargas JL, Di Polo A. The molecular basis of retinal ganglion cell death in glaucoma. Prog Retin Eye Res. 2012;31(2):152–181.

17. Livne-Bar I, Lam S, Chan D, et al. Pharmacologic inhibition of reactive gliosis blocks TNF-α-mediated neuronal apoptosis. Cell Death Dis. 2016;7(9):e2386.

18. Cooper ML, Collyer JW, Calkins DJ. Astrocyte remodeling without gliosis precedes optic nerve Axonopathy. Acta Neuropathol Commun. 2018;6(1):38.

19. Tehrani S, Johnson EC, Cepurna WO, Morrison JC. Astrocyte processes label for filamentous actin and reorient early within the optic nerve head in a rat glaucoma model. Invest Ophthalmol Vis Sci. 2014;55(10):6945–6952.

20. Lopez NN, Clark AF, Tovar-Vidales T. Isolation and characterization of human optic nerve head astrocytes and lamina cribrosa cells. Exp Eye Res. 2020;197:108103.

21. Yang P, Hernandez MR. Purification of astrocytes from adult human optic nerve heads by immunopanning. Brain Res Brain Res Protoc. 2003;12(2):67–76.

22. Tezel G, Hernandez MR, Wax MB. In vitro evaluation of reactive astrocyte migration, a component of tissue remodeling in glaucomatous optic nerve head. Glia. 2001;34(3):178–189.

23. Hariani HN, Ghosh AK, Rosen SM, et al. Lysyl oxidase like-1 deficiency in optic nerve head astrocytes elicits reactive astrocytosis and alters functional effects of astrocyte derived exosomes. Exp Eye Res. 2024;240:109813.

24. Urban Z, Agapova O, Hucthagowder V, Yang P, Starcher BC, Hernandez MR. Population differences in elastin maturation in optic nerve head tissue and astrocytes. Invest Ophthalmol Vis Sci. 2007;48(7):3209–3215.

25. Caliari SR, Burdick JA. A practical guide to hydrogels for cell culture. Nat Methods. 2016;13(5):405–414.

26. Puschmann TB, Zandén C, De Pablo Y, et al. Bioactive 3D cell culture system minimizes cellular stress and maintains the in vivo-like morphological complexity of astroglial cells. Glia. 2013;61(3):432–440.

27. Liu B, McNally S, Kilpatrick JI, Jarvis SP, O’Brien CJ. Aging and ocular tissue stiffness in glaucoma. Surv Ophthalmol. 2018;63(1):56–74.

28. Lee PY, Yang B, Hua Y, et al. Real-time imaging of optic nerve head collagen microstructure and biomechanics using instant polarized light microscopy. Exp Eye Res. 2022;217:108967.

29. Alarcon-Martinez L, Shiga Y, Villafranca-Baughman D, et al. Pericyte dysfunction and loss of interpericyte tunneling nanotubes promote neurovascular deficits in glaucoma. Proc Natl Acad Sci U S A. 2022;119(7). doi:10.1073/pnas.2110329119

30. Wareham LK, Calkins DJ. The Neurovascular Unit in Glaucomatous Neurodegeneration. Front Cell Dev Biol. 2020;8:452.

31. Alarcon-Martinez L, Shiga Y, Villafranca-Baughman D, et al. Neurovascular dysfunction in glaucoma. Prog Retin Eye Res. 2023;97:101217.

32. Elkington AR, Inman CB, Steart PV, Weller RO. The structure of the lamina cribrosa of the human eye: an immunocytochemical and electron microscopical study. Eye. 1990;4 (Pt 1):42–57.

33. Reynaud J, Lockwood H, Gardiner SK, Williams G, Yang H, Burgoyne CF. Lamina Cribrosa Microarchitecture in Monkey Early Experimental Glaucoma: Global Change. Invest Ophthalmol Vis Sci. 2016;57(7):3451–3469.

34. Brooks DE, Arellano E, Kubilis PS, Komaromy AM. Histomorphometry of the porcine scleral lamina cribrosa surface. Vet Ophthalmol. 1998;1(2-3):129–135.

35. Prasanna G, Krishnamoorthy R, Clark AF, Wordinger RJ, Yorio T. Human optic nerve head astrocytes as a target for endothelin-1. Invest Ophthalmol Vis Sci. 2002;43(8):2704–2713.

36. Johnson EC, Deppmeier LM, Wentzien SK, Hsu I, Morrison JC. Chronology of optic nerve head and retinal responses to elevated intraocular pressure. Invest Ophthalmol Vis Sci. 2000;41(2):431–442.

37. Oberheim NA, Takano T, Han X, et al. Uniquely hominid features of adult human astrocytes. J Neurosci. 2009;29(10):3276–3287.

38. Sun D, Lye-Barthel M, Masland RH, Jakobs TC. The morphology and spatial arrangement of astrocytes in the optic nerve head of the mouse. J Comp Neurol. 2009;516(1):1–19.

39. Waxman S, Brazile BL, Yang B, et al. Lamina cribrosa vessel and collagen beam networks are distinct. Exp Eye Res. 2022;215:108916.

40. Waxman S, Quinn M, Donahue C, et al. Individual astrocyte morphology in the collagenous lamina cribrosa revealed by multicolor DiOlistic labeling. Exp Eye Res. 2023;230:109458.

41. Sun D, Jakobs TC. Structural remodeling of astrocytes in the injured CNS. Neuroscientist. 2012;18(6):567–588.

42. Luna G, Keeley PW, Reese BE, Linberg KA, Lewis GP, Fisher SK. Astrocyte structural reactivity and plasticity in models of retinal detachment. Exp Eye Res. 2016;150:4–21.

43. Bushong EA, Martone ME, Jones YZ, Ellisman MH. Protoplasmic astrocytes in CA1 stratum radiatum occupy separate anatomical domains. J Neurosci. 2002;22(1):183–192.

44. Stokum JA, Kwon MS, Woo SK, et al. SUR1-TRPM4 and AQP4 form a heteromultimeric complex that amplifies ion/water osmotic coupling and drives astrocyte swelling. Glia. 2018;66(1):108–125.

45. Escartin C, Galea E, Lakatos A, et al. Reactive astrocyte nomenclature, definitions, and future directions. Nat Neurosci. 2021;24(3):312–325.

46. Gan WB, Grutzendler J, Wong WT, Wong RO, Lichtman JW. Multicolor “DiOlistic” labeling of the nervous system using lipophilic dye combinations. Neuron. 2000;27(2):219–225.

47. Nolte C, Matyash M, Pivneva T, et al. GFAP promoter-controlled EGFP-expressing transgenic mice: a tool to visualize astrocytes and astrogliosis in living brain tissue. Glia. 2001;33(1):72–86.

48. Virtanen P, Gommers R, Oliphant TE, et al. SciPy 1.0: fundamental algorithms for scientific computing in Python. Nat Methods. 2020;17(3):261–272.

49. Hunter JD. Matplotlib: A 2D Graphics Environment. Comput Sci Eng. May-June 2007;9(3):90–95.

50. Waskom M. seaborn: statistical data visualization. J Open Source Softw. 2021;6(60):3021.

51. Jan NJ, Lathrop K, Sigal IA. Collagen Architecture of the Posterior Pole: High-Resolution Wide Field of View Visualization and Analysis Using Polarized Light Microscopy. Invest Ophthalmol Vis Sci. 2017;58(2):735–744.

52. Gogola A, Jan NJ, Lathrop KL, Sigal IA. Radial and Circumferential Collagen Fibers Are a Feature of the Peripapillary Sclera of Human, Monkey, Pig, Cow, Goat, and Sheep. Invest Ophthalmol Vis Sci. 2018;59(12):4763–4774.

53. Lee PY, Schilpp H, Naylor N, Watkins SC, Yang B, Sigal IA. Instant polarized light microscopy pi (IPOLπ) for quantitative imaging of collagen architecture and dynamics in ocular tissues. Opt Lasers Eng. 2023;166. doi:10.1016/j.optlaseng.2023.107594

54. Ogata K, Kosaka T. Structural and quantitative analysis of astrocytes in the mouse hippocampus. Neuroscience. 2002;113(1):221–233.

55. May CA, Lütjen-Drecoll E. Morphology of the murine optic nerve. Invest Ophthalmol Vis Sci. 2002;43(7):2206–2212.

56. Kotecha A, Izadi S, Jeffery G. Age-related changes in the thickness of the human lamina cribrosa. Br J Ophthalmol. 2006;90(12):1531–1534.

57. Quigley HA, Brown AE, Morrison JD, Drance SM. The size and shape of the optic disc in normal human eyes. Arch Ophthalmol. 1990;108(1):51–57.

58. Brazile BL, Hua Y, Jan NJ, Wallace J, Gogola A, Sigal IA. Thin Lamina Cribrosa Beams Have Different Collagen Microstructure Than Thick Beams. Invest Ophthalmol Vis Sci. 2018;59(11):4653–4661.

59. Nadler Z, Wang B, Schuman JS, et al. In vivo three-dimensional characterization of the healthy human lamina cribrosa with adaptive optics spectral-domain optical coherence tomography. Invest Ophthalmol Vis Sci. 2014;55(10):6459–6466.

60. Wang B, Lucy KA, Schuman JS, et al. Decreased Lamina Cribrosa Beam Thickness and Pore Diameter Relative to Distance From the Central Retinal Vessel Trunk. Invest Ophthalmol Vis Sci. 2016;57(7):3088–3092.

61. Coudrillier B, Geraldes DM, Vo NT, et al. Phase-Contrast Micro-Computed Tomography Measurements of the Intraocular Pressure-Induced Deformation of the Porcine Lamina Cribrosa. IEEE Trans Med Imaging. 2016;35(4):988–999.

62. Jan NJ, Gomez C, Moed S, et al. Microstructural Crimp of the Lamina Cribrosa and Peripapillary Sclera Collagen Fibers. Invest Ophthalmol Vis Sci. 2017;58(9):3378–3388.

63. Voorhees AP, Jan NJ, Sigal IA. Effects of collagen microstructure and material properties on the deformation of the neural tissues of the lamina cribrosa. Acta Biomater. 2017;58:278–290.

64. Schlamp CL, Li Y, Dietz JA, Janssen KT, Nickells RW. Progressive ganglion cell loss and optic nerve degeneration in DBA/2J mice is variable and asymmetric. BMC Neurosci. 2006;7:66.

65. Guttenplan KA, Stafford BK, El-Danaf RN, et al. Neurotoxic Reactive Astrocytes Drive Neuronal Death after Retinal Injury. Cell Rep. 2020;31(12):107776.

66. Li J, Pan L, Pembroke WG, et al. Conservation and divergence of vulnerability and responses to stressors between human and mouse astrocytes. Nat Commun. 2021;12(1):3958.

67. Vicic N, Guo X, Chan D, Flanagan JG, Sigal IA, Sivak JM. Evidence of an Annexin A4 mediated plasma membrane repair response to biomechanical strain associated with glaucoma pathogenesis. J Cell Physiol. 2022;237(9):3687–3702.

68. Murphy R, Irnaten M, Hopkins A, et al. Matrix Mechanotransduction via Yes-Associated Protein in Human Lamina Cribrosa Cells in Glaucoma. Invest Ophthalmol Vis Sci. 2022;63(1):16.

69. Morgan JE. Optic nerve head structure in glaucoma: astrocytes as mediators of axonal damage. Eye. 2000;14 (Pt 3B):437–444.

70. Jan NJ, Sigal IA. Collagen fiber recruitment: A microstructural basis for the nonlinear response of the posterior pole of the eye to increases in intraocular pressure. Acta Biomater. 2018;72:295–305.

71. Lee PY, Fryc G, Gnalian J, et al. Direct measurements of collagen fiber recruitment in the posterior pole of the eye. Acta Biomater. 2024;173:135–147.

72. Ling YTT, Pease ME, Jefferys JL, Kimball EC, Quigley HA, Nguyen TD. Pressure-Induced Changes in Astrocyte GFAP, Actin, and Nuclear Morphology in Mouse Optic Nerve. Invest Ophthalmol Vis Sci. 2020;61(11):14.

73. Gallego BI, Salazar JJ, de Hoz R, et al. IOP induces upregulation of GFAP and MHC-II and microglia reactivity in mice retina contralateral to experimental glaucoma. J Neuroinflammation. 2012;9:92.

74. Dillinger AE, Weber GR, Mayer M, et al. CCN2/CTGF—A Modulator of the Optic Nerve Head Astrocyte. Frontiers in Cell and Developmental Biology. 2022;10. doi:10.3389/fcell.2022.864433

